# Crystal Structure of the Dog Allergen Can f 6

**DOI:** 10.1101/551549

**Authors:** Gina M Clayton, Janice White, John W Kappler, Sanny K Chan

## Abstract

Lipocalins represent the most important protein family of the mammalian respiratory allergens. Four of the seven named dog allergens are lipocalins: Can f 1, Can f 2, Can f 4, and Can f 6. We present the structure of Can f 6 along with data on the biophysical and biological activity of this protein in comparison with other animal lipocalins. The Can f 6 structure displays the classic lipocalin calyx-shaped ligand binding cavity within a central β-barrel similar to other lipocalins. Despite low sequence identity between the different dog lipocalin proteins, there is a high degree of structural similarity. On the other hand, Can f 6 has a similar primary sequence to cat, horse, mouse lipocalins as well as a structure that may underlie their cross reactivity. Interestingly, the entrance to the ligand binding pocket is capped by a His instead of the usually seen Tyr that may help select its natural ligand binding partner. Our highly pure recombinant Can f 6 is able to bind to human IgE (hIgE) demonstrating biological antigenicity.

## Introduction

Analyses over the last 2 decades have revealed that most allergens can be grouped into a few families of proteins(1). One such family is made up of lipocalins. These are amongst the most important inhalant animal allergens. As far as their functions in their host of origin are concerned, these, depending on the lipocalin involved, include a number of properties such as pheromone transport, prostaglandin synthesis, retinoid binding, odorant binding and cancer cell interactions(1). Lipocalins are small proteins, usually composed of less than 200 amino acids. In spite of their shared ability to act as allergens they have limited sequence homology with each other, even as low as 20%. Yet for the lipocalins whose structures have been solved, they have a common tertiary structure consisting of an 8 stranded antiparallel beta barrel with a hydrophobic ligand binding pocket(2). Human sensitization has been identified against lipocalins from cows, guinea pigs, horses, cats, mice, rats(3) and several arthropods(4). Human sensitization to dogs and cats is a known major risk factor for the development of asthma and allergic rhinitis. Because the prevalence of allergic disease to furry animals unfortunately continues to increase worldwide(5–7) it will be useful to expand our knowledge of the structure and function of the lipocalins that are most often involved in such human diseases.

Four of the 7 named dog allergens, Can f 1, Can f 2, Can f 4, and Can f 6, are lipocalins, though their ligands and biological functions are poorly defined. One study of patients who were sensitized to unfractionated dog dander showed that variable numbers of the patients had IgE binding the different dog allergens, ranging from 64% allergic to Can f 1 to 27% allergic to Can f 3(8). Symptoms from patients when exposed to dogs range from those of asthma to no symptoms at all, with percent responses depending to some extent on the dog allergens to which the sensitized individuals were exposed(9).

In the studies described here we chose to focus on one of the dog lipocalin allergens, Can f 6. Recent reports suggests that Can f 6 is a key lipocalin responsible for allergic symptoms(10). Responses to Can f 6 are present in between 38%(11) and 56.3%(12) of dog sensitized individuals and Can f 6 allergic individuals are more likely to suffer from allergic rhinitis symptoms at a rate of 86% vs non-sensitized at 53% and those with asthma symptoms are found to be sensitized at 64% vs 38% in non-sensitized(10). Therefore, Can f 6 is a major allergen driving disease amongst those people who are allergic to dogs. Moreover, in some individuals, antibodies to the cat lipocalin, Fel d 4, and/or the horse lipocalin, Equ c 1, cross react with Can f 6 suggesting some structural and sequence similarities between the 3 lipocalins(11,13). Since mouse Mus m 1 cross reacts with Equ c 1(14), it is likely that there is also cross reactivity with Can f 6. Although the structure for Fel d 4 is unknown, the ones for Equ c 1(15) and Mus m 1(16) are, so a comparison of the structures of Equ c 1 and Mus m 1 with Can f 6 might reveal the reasons for their immunological cross reactivity.

Here the structure of the clinically relevant Can f 6 allergen was determined at 2 Å (Table 1) resolution along with human antibody binding data that shows its biologic activity. Although structurally similar to other dog lipocalins, Can f 6’s primary sequence is more similar to other mammalian lipocalins. Unique regions in the capping region and shape of the ligand binding pocket may play a role in ligand recognition, binding, and ultimate function in vivo. We also describe both unique and common features of Can f 6 and similar lipocalins that underlie their allergencity.

**Table 1.**
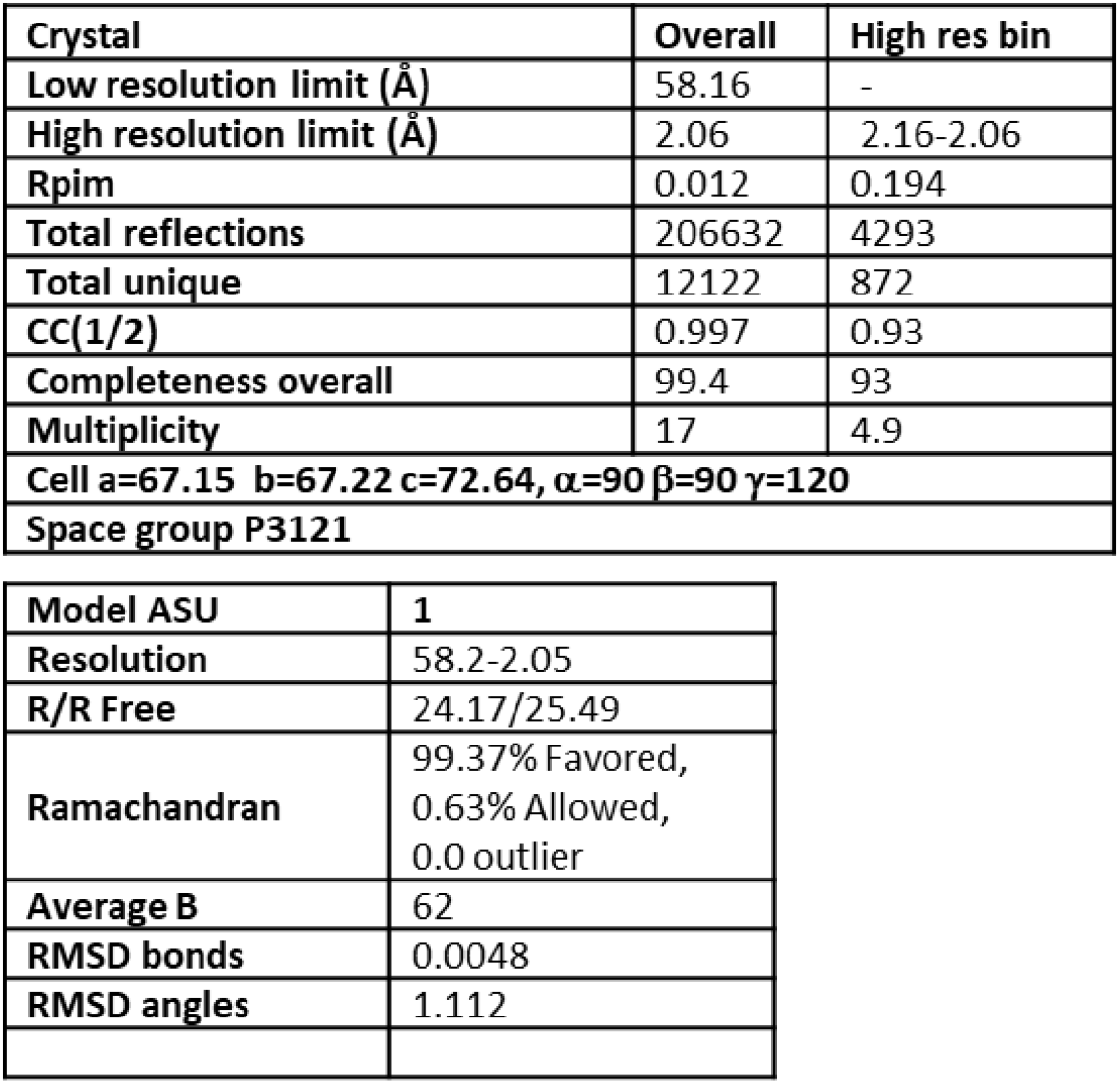
Crystallographic data and model refinement statistics for Can f 6.

| Crystal | Overall | High res bin |
| --- | --- | --- |
| Low resolution limit ($\text{\AA}$) | 58.16 | - |
| High resolution limit ($\text{\AA}$) | 2.06 | 2.16-2.06 |
| R<sub>p</sub>im | 0.012 | 0.194 |
| Total reflections | 206632 | 4293 |
| Total unique | 12122 | 872 |
| CC(1/2) | 0.997 | 0.93 |
| Completeness overall | 99.4 | 93 |
| Multiplicity | 17 | 4.9 |
| Cell a=67.15 b=67.22 c=72.64, $\alpha=90^\circ$ $\beta=90^\circ$ $\gamma=120^\circ$ | | |
| Space group P3121 |  |  |

|  |  |
| --- | --- |
| Model ASU | 1 |
| Resolution | 58.2-2.05 |
| R/R Free | 24.17/25.49 |
| Ramachandran | 99.37% Favored,<br>0.63% Allowed,<br>0.0 outlier |
| Average B | 62 |
| RMSD bonds | 0.0048 |
| RMSD angles | 1.112 |

## Material and Methods

### Cloning, production, purification, and biochemical characterization

The cDNA sequence of *Can f 6*(11) was cloned with a C-terminal His_6_-tag and expressed in the periplasmic space of the Rosetta strain of *E. coli*, as previously described, with minor modifications(17). Can f 6 was purified using a HisPur Ni-NTA Superflow Agarose nitrilotriacetic acid (NTA) column, followed by size exclusion on a Superdex 10/300 column (GE), and anion exchange chromatography (GE). The purified protein, Can f 6, was confirmed by Matrix Assisted Laser Desorption Ionization time-of-flight/mass spectrometer (MALDI TOF/MS) using the ThermoScientific LTQ Orbitrap Velos Pro mass spectrometer.

Measurements of the thermal stability of Can f 6 were made using an Agilent Stratagene MX3005P qPCR thermal shift assay as previously described(17). Briefly, the fluorescence of Sypro Orange was monitored as temperatures were increased, leading to unfolding of the Can f 6 protein.

### Crystallization

Purified recombinant Can f 6 at 15mg/ml was buffer exchanged into 10mM Tris pH 8 and 100mM NaCl. Initial crystallization trials were undertaken at room temperature using the mosquito LCP Nanoliter Protein crystallization robot. Crystals were grown and further optimized by the hanging drop vapor diffusion method containing a reservoir solution of 6% v/v 2-propanol, 0.1M sodium acetate trihydrate pH 4.5, and 26% v/v polyethylene glycol monomethyl ether 550.

### Data collection, structure determination and refinement

Can f 6 crystals were harvested and cryoprotected in a reservoir solution containing 35% PEG 500 MME followed by storage in liquid N_2_. Data were collected on a single crystal at the NE-CAT 24-ID-C beam line Advanced Photon Source, Argonne National Laboratory. Data were processed in XDS Aimless and further analyzed in Phenix Xtriage(18).

### Structural determination

The structure of Can f 6 was determined by molecular replacement in Phaser(19) using Can f 4 (PDB ID: 4ODD)(20) as an initial search model. The resulting solution was further modelled and refined in the CCP4(21) suite of programs using Coot and Refmac. Phenix Validation and Refine(18) were used to confirm the final structure. The crystal packing of Can f 6 was analyzed in PISA(22) and its secondary structure was analyzed in STRIDE(23). See Table 1 for data and refinement statistics. Pymol was used for analysis of the structure and to generate figures(24). The atomic coordinates and the structure factors have been deposited in the RCSB Protein Data Bank with accession code 6NRE.

The amino acid residues of the ligand-binding sites and the volumes of the central cavities were determined using CASTp(25) and Voidoo(26) was used to generate a cavity volume map and visualized as Connolly surfaces(27). Post-translational modifications were analyzed with NetNGlyc 1.0 server(28) for potential N-glycosylated sites, NetOGlyc 4.0 server was used to predict GalNAc O-glycosylation sites(29), NetPhos 3.1 Server(30) was used to predicted phosphorylation sites, and Sulfinator(31) was used to predict tyrosine sulfation.

Sequence alignments for Can f 6, Can f 2, Can f 4, Fel d 4, Equ c 1, and Mus m 1 was performed in Clustal Omega(32). Superpositions of Can f 6, Can f 2 (PDB ID: 3L4R), Can f 4 (PDB ID: 4ODD), Equ c 1 (PDB ID: 1EW3), and Mus m 1 (PDB ID: 1QY0) were performed using Coot and Pymol.

### Detection of Human Antibody Binding by ELISA

Patients’ sample data were downloaded from the National Jewish Health Research Database, Data Set Identifier: 473-06-20-2018. Human serum samples banked by National Jewish Health under IRB approved protocol were retrieved after IRB approval. Sera from 5 patients who had dog IgE >5 kU/L (ImmunoCAP; Phadia, Uppsala Sweden) and skin prick testing >3×3mm to AP dog extract were selected at random for analysis. Sera from 5 age matched controls who had IgE<0.35 kU/L and negative skin testing were then selected from the biobank. For detection of biological activity by ELISA, plates were coated with recombinant Can f 6 at 50 μg/ml overnight at 4°C and then incubated with serially diluted sera. Bound IgE was detected using alkaline phosphatase-conjugated goat anti-human secondary antibodies (A18796, Invitrogen) followed by incubation with p-nitrophenyl phosphate and measurement by spectrophotometry. The average baseline background OD405nm was subtracted from all samples and readings from the 1:10 dilution was used for analysis of significance using an unpaired t-test.

## Results and Discussion

### Overall structure of Can f 6

The recombinant Can f 6 crystallized in space group P3_1_2_1_ with one monomer in the asymmetric unit (ASU) and was refined to a resolution of 2.06 Å. Final refinement and model statistics are in Table 1. The overall architecture of Can f 6 contains a highly conserved lipocalin fold of an 8 stranded anti-parallel β-barrel enclosing a central ligand binding cavity core, a four-turn C-terminal α-helix that packs against the β-barrel, and a short 9th β-strand (β_I_) at the C-terminus (Fig. 1A). The top open end of the β-barrel, which marks the entrance to the presumptive ligand-binding cavity, is comprised of four β-hairpin turns. The other side of the β-barrel is closed by one β-hairpin turn, two smaller β-hairpin turns, and an N-terminal coil (Fig. 1A). There is a disulfide bond between residues Cys69 and Cys162 that connects the C-terminal part of the protein to the central β-barrel. Three lipocalin Structurally Conserved Regions (SCRs) are present (Fig. 1A and B) that contain conserved residues. The SCRs hold together the N- and C-terminus of the overall fold (Fig. 1A) under the ligand cavity calyx. By comparison, the cavity cap and calyx are more distinct in sequence and conformation when considered across all lipocalins. For example, in the structures of the lipocalins apo, holo Retinol-binding protein (RBP) (33) and apolioprotein M (34), the ligand cap conformations are different and have a more open architecture than the ligand cavity cap described here.

**Fig 1.**
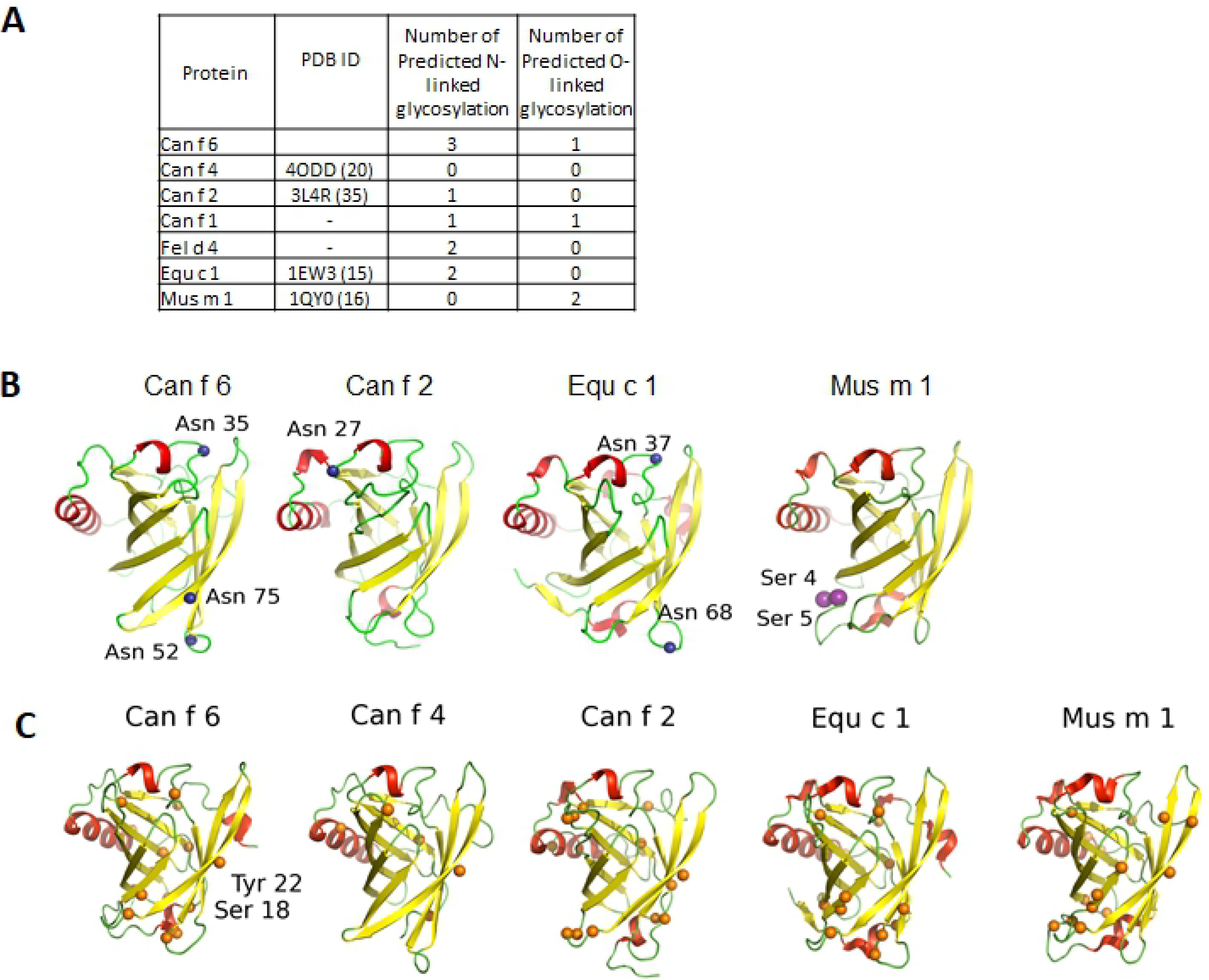
**(A)** The overall architecture of Can f 6 displays the classic lipocalin structure. Eight antiparallel β-strands (yellow) form a central ligand binding cavity (β _A-I_) flanked by C-terminal α-helix (posteriorly located in the figure) and three 3_10_ -helices (red), with the connecting loops (L, in green) numbered. A disulfide bond (**DS**, cyan) clamps the C-terminal loop to the beta core via β_D_. The cavity opening is covered by L1 and a loop-3_10_ helix-loop that links together β _A_ and β_D_. Three core lipocalin Structurally Conserved Regions (**SCR**) that hold the overall fold together are labeled: **SCR1** connects the N-terminal loop to the C-terminal **SCR3** which also interacts with **SCR2**. (**B**) Figure A is rotated by 90° out of the plane and shows the main residues of the core SCR. In **SCR3**, the side chain and distal guanidium group of a conserved Arg125 are stabilized by hydrogen bonds (dashed lines) to the backbone carbonyls of other SCR residues and a Cation-Pi interaction with a conserved **SCR1** aromatic residue, here Trp21. Other residues in the SCR also form numerous non-covalent interactions. **(C)** Purified Can f 6 was assayed by Thermofluor shift to demonstrate the stability of our recombinant Can f 6. The Tm is 67.9°C with standard deviation of 1.7°C

Can f 6 crystalized as a monomer reflecting its behavior during purification and on size exclusion chromatography at pH 8. The protein is quite stable to heat (Fig.1C) which is perhaps helped by the disulfide bond and the interactions of the SCR. PISA analysis of the crystal lattice packing around the Can f 6 monomer reveals 3 adjacent lattice packing molecules. The largest packing surface, at symmetry operator position Y, X, -Z, forms a dimer buried surface area of 736 Å^2^ (Table 2). The dimer interface is the same as one of the dimers observed in the other deposited structure of Can f 6 (PDB ID: 5X7Y) which forms a tetramer in the ASU.

**Table 2.** The Can f 6 monomer-monomer surface interlaces within the crystal packing generated in PISA.

| Monomer-monomer interface |  |  |  |  |
| --- | --- | --- | --- | --- |
| Surface residues | 148 (92%) |  |  |  |
| Symmetry Operation | y, x, -z | -y+1, x-y+1, z+1/3 | -x, -x+y, -z+1/3 | x-y, -y+1, -z-1/3 |
| Interface residues | 27 (17%) | 17 (10%) | 16 | 4 (3%) |
| Buried ASA (Å<sup>2</sup>) | 736 (9.4%) | 510 | 420 (5%) | 69 (1%) |
| Hydrogen bonds | 14 | 2 | 0 | 1 |
| Salt bidges | 0 | 0 | 0 | 0 |
| Complex Formation Significance Score (CSS) | 0.1 | 0 | 0 | 0 |
| Total ASA | 7826 |  |  |  |

Interface surfaces of less than 1000 Å^2^ between monomers can be classified as having weak transient interactions(35). The PISA analysis of the Can f 6 crystal did not reveal any specific interactions that might result in the formation of a stable quaternary structure in solution(22). The dimeric interface of the Can f 6 structure is considered below that for the complexation criteria. However, the dimeric crystal packing interface consists of 27 residues and 14 hydrogen bonds from each monomer (Table 2) and as such would be susceptible to changes in pH conditions during purification and crystallization.

To date, lipocalin structures reveal buried interface surfaces that vary considerably from 390-23475 Å^2^(35) as dimers, trimers, or tetramers without a consistent quaternary pattern. The crystal lattice packing of Can f 6 does not resemble those observed in the structures of Can f 4(20) nor Equ c 1(15). In the Can f 4 structure there is a dimer interface (largest interface 640 Å^2^)(20). Similarly, in the Equ c 1 structure there is also a dimer interface of (interface 1070 Å^2^)(15). In the structure of Mus m 1 the dimer interface is much smaller (interface of 389 Å^2^)(35).

The packing interfaces in the lipocalin structures may be physiologically relevant or may simply be a consequence of crystallization conditions. Differences in the monomeric and transient multimeric states of lipocalins are theorized to contribute to a protein’s allergenicity and are likely unique to each allergen(4). For example, the dimerization of the lipocalin allergen Bet v 1 has been demonstrated to increase allergenicity in skin tests of allergic mice compared to monomers(36). However, the role of multimerization in allergenic lipocalins is inconsistent. It is not known if Can f 6 forms multimers in solution at different pH conditions as reported for other lipocalins(35).

### Comparison of the structure and sequence of Can f 6 with those of other mammalian lipocalins

Some members of the lipocalin family have limited sequence homology with each other. For example, of the known dog lipocalins, Can f 2 is only 21% identical to Can f 4 and likewise Can f 6 is 25% identical to Can f 2 and 29% identical to Can f 4 (Fig. 2A and 3). On the other hand, Can f 6 is 66%; 57%; and 47% identical to the cat lipocalin, Fel d 4; the major horse allergen, Equ c 1; and the mouse lipocalin Mus m 1 respectively (Fig. 2B and 3). Cross reactivity with IgE antibody binding has been show between Can f 6, Fel d 4 and Equ c 1(11) as well as between Equ c 1 and Mus m 1 (14) which would also suggest that Can f 6 and Mus m 1 would also cross react.

**Fig. 2.**
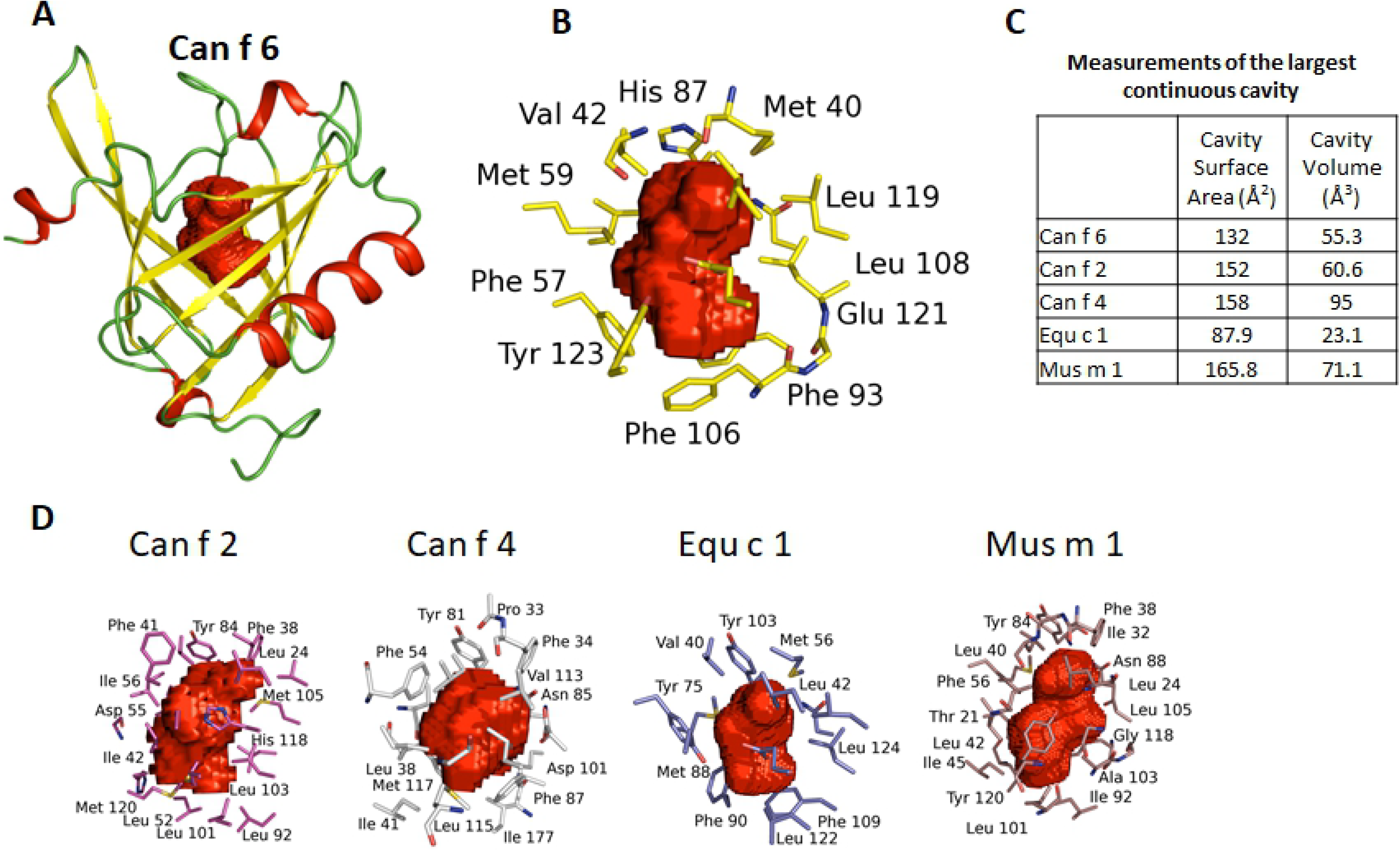
(A). Primary sequence alignment of lipocalins. The secondary structure for Can f 6 is drawn above the sequences. The colors correspond to the model of Can f 6 depicted in Fig. 1 (A). Despite their conserved lipocalin architecture there is low sequence similarity between the dog lipocalins Can f 1, 2, 4, and Can f 6. The only disulfide bond (C) occurs between C67 and C160. **(B)** Primary sequence alignment of the Can f 6 dog lipocalin, cat (Fel d 4), horse (Equ c 1), and mouse (Mus m 1). There is much more sequence similarity in these lipocalins compared to those of the dog lipocalins. Alignment residues are as: “*” Identical amino acid, “:” Similar amino acid, and “.” Slightly similar amino acid (30). 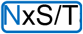are predicted N glycosylation sites (28), 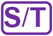are predicted O glycosylation sites (29), <u>S/T/T</u> are predicted phosphorylation sites (30), and 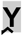 are predicted sulfation sites (31).

**Fig. 3.**
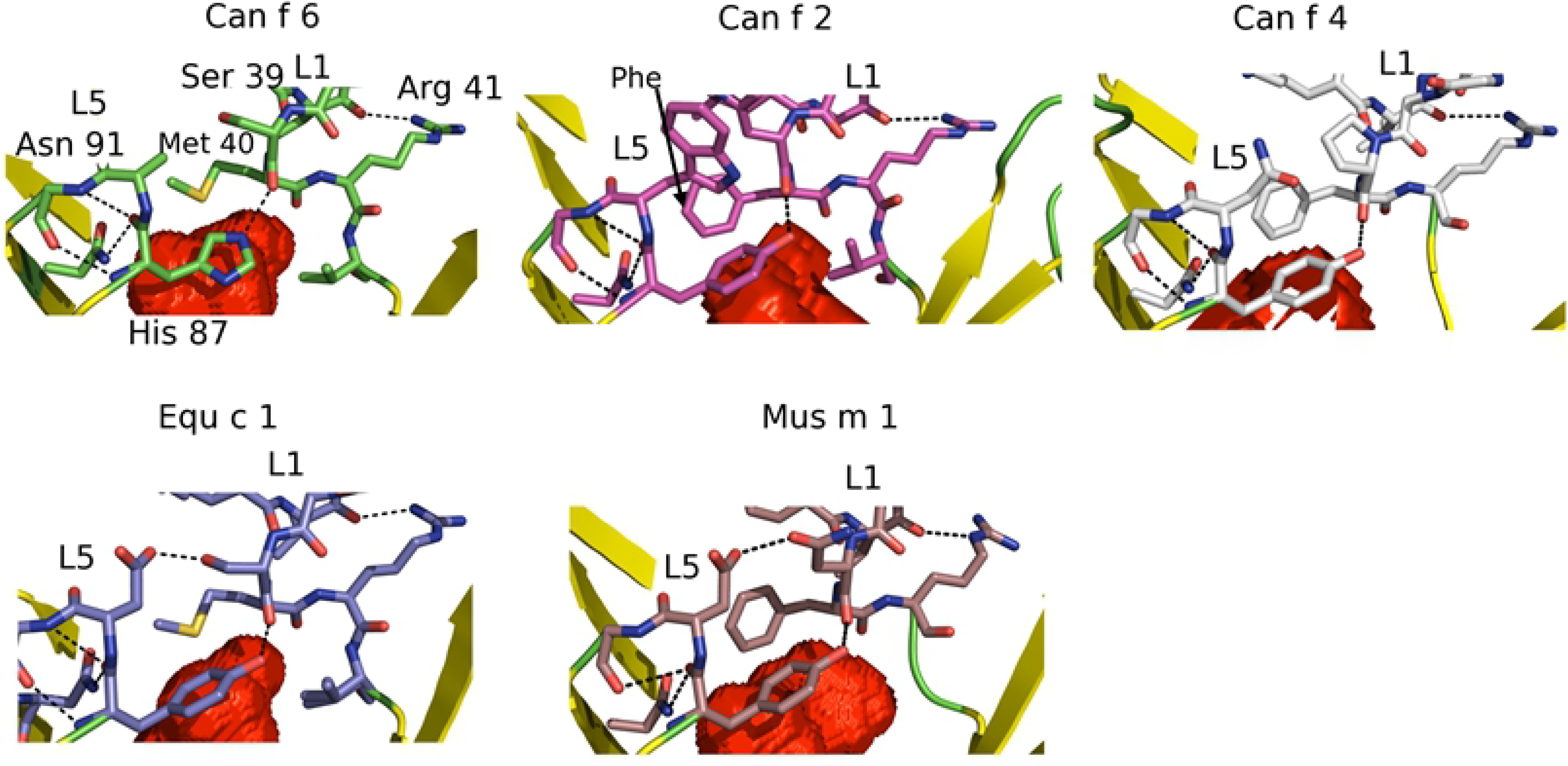
The sequence alignment of lipocalins threaded onto the structure of Can f 6. The sequence homology between the dog lipocalin Can f 6, 4, and 2 are more distinct compared to that between Can f 6 and Equ c 1, Fel d 4, and Mus m 1.

The crystal structures of Can f 2 (PDB ID: 3L4R)(37), Can f 4 (PDB ID: 4ODD)(20), Equ c 1 (PDB ID: 1EW3)(15), and Mus m 1 (PDB ID: 1QY0)(16) have been determined previously. Currently, no structure of Fel d 4 has been reported. Superposition of the structures of Can f 2, Can f 4, and Equ c 1 onto Can f 6 yields an rms deviation for Cα backbone atoms of 1.32, 1.48, and 1.38 Å respectively. The overall topology is similar for all 4 (Fig. 4). The main structural differences are the β-strand turns and the folds preceding the C-terminal helices. For example, secondary structure analysis (23) of the lipocalins here, reveals that only Can f 2 has a short 3_10_ helix (res 124-126) preceding the C-terminal α-helix (Fig. 2) whereas the differences in secondary structures of the other lipocalins likely interfere with 3_10_ helix formation. In Can f 6, Can f 4, and Equ c 1, the equivalent region is a turn towards the C-terminal α-helix. In both Can f 6 and Equ c 1, the turn has a Pro in β_F_ (127 in Can f 6 and 143 in Equ c 1) that stacks against a Tyr in β_A_ (22 in Can f 6 and 38 in Equ c 1). Pro127 stabilizes this turn via a backbone carbonyl hydrogen bond to the side chain of Arg159 (Arg175 in Equ c 1). By comparison Can f 4, has a β_F_ Gly120 and Ala121 that packs against the β_A_ Lys16. However, it is unclear if these conformational differences are normally occurring or due to crystal packing observed in the lattice.

**Fig. 4.**
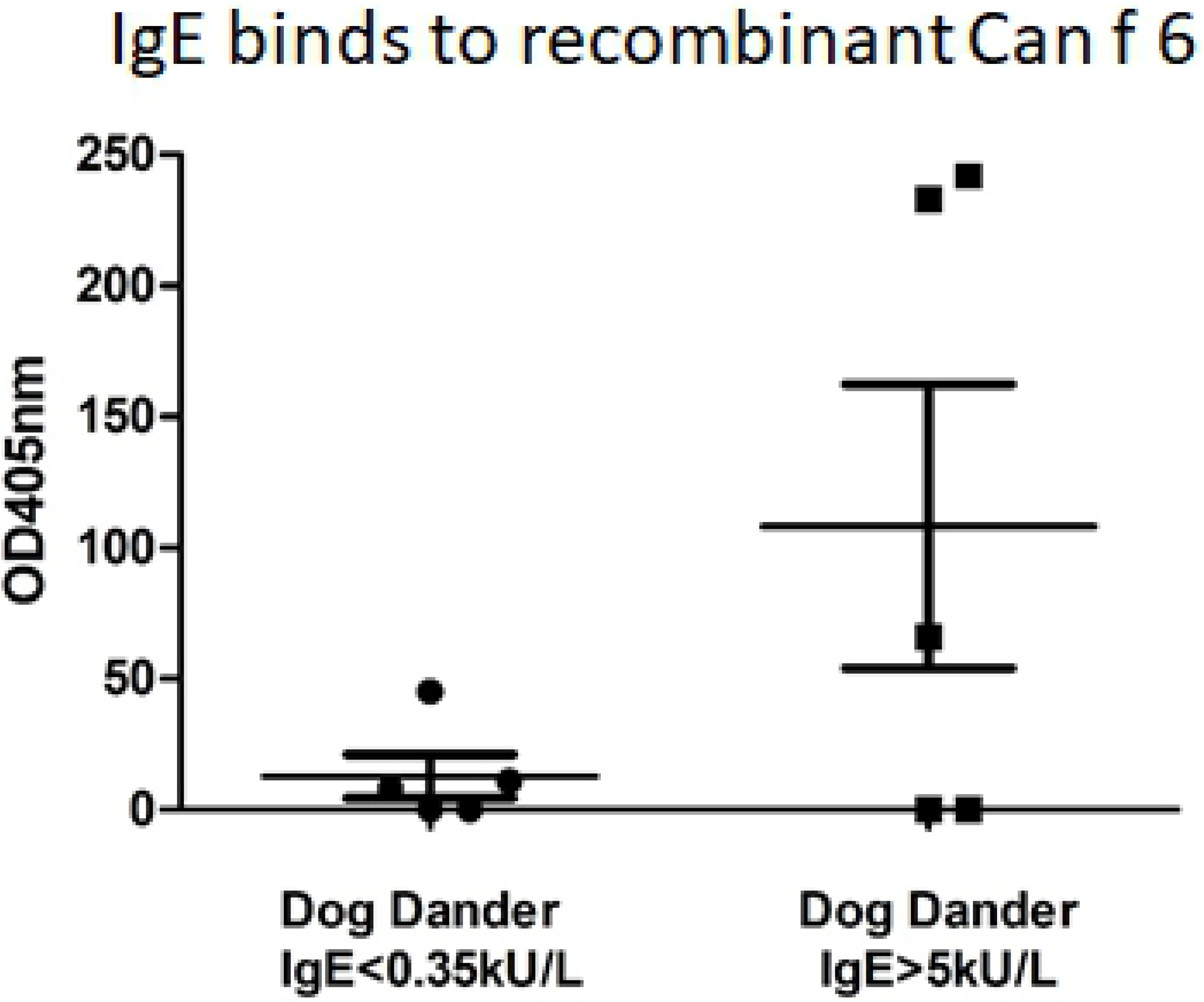
Structural overlay of Can f 6 (yellow) on Can f 2 (red) and Can f 4 (grey) Equ c 1 (purple), and Mus m 1 (brown). The overall architecture of lipocalins is highly conserved with a β-barrel that encloses a hydrophobic core. The major differences are outside the core.

Although lipocalins are of similar size (Table 3 and Fig. 2), the residue differences mean they do not necessarily have a similar overall charge, immunogenic sites, nor post translational modification patterns. Predicted post translational modifications are shown (Fig. 2 and 5). The predicted N-linked glycosylation pattern for Can f 6 is different when compared to the other dog lipocalins here (Fig. 2 and 5). Two of the predicted N-linked glycosylation sites (38) of the immunologically cross-reactive Can f 6, Fel d 4, and Equ c 1 are at the similar sites, but are not present in Mus m 1 (Fig. 5B). Recombinant Can f 6 was expressed in E. Coli and was not phosphorylated as determined by mass spec (data not shown). However, the prediction algorithm for Can f 6 suggests numerous potential sites (Fig. 2 and 5C).

**Table 3.** Molecular masses, numbers of positively and negatively charged amino acids and theoretical pl of Can f 6, Can f 4, and Can f2 using ProtParam.

| Protein | Total Amino Acids | Molecular Mass (kDa) | Positively charged amino acids (Arg + Lys) | Negatively charged amino acids (Asp + Glu) | pI | Resolution (Å) |
| --- | --- | --- | --- | --- | --- | --- |
| Can f 6 | 179 | 20667 | 21 | 34 | 4.75 | 2 (Now) |
| Can f 4 | 159 | 17693 | 19 | 20 | 6.17 | 2.6 (20) |
| Can f 2 | 163 | 18448 | 14 | 27 | 4.82 | 1.45 (32) |
| Can f 1 | 156 | 17319 | 18 | 21 | 5.66 |  |
| Fel d 4 | 170 | 19573 | 19 | 31 | 4.76 |  |
| Equ c 1 | 172 | 20097 | 20 | 34 | 4.51 | 2.3 (15) |
| Mus m 1 | 163 | 18765 | 19 | 31 | 4.84 | 1.8 (16) |

**Fig. 5.**
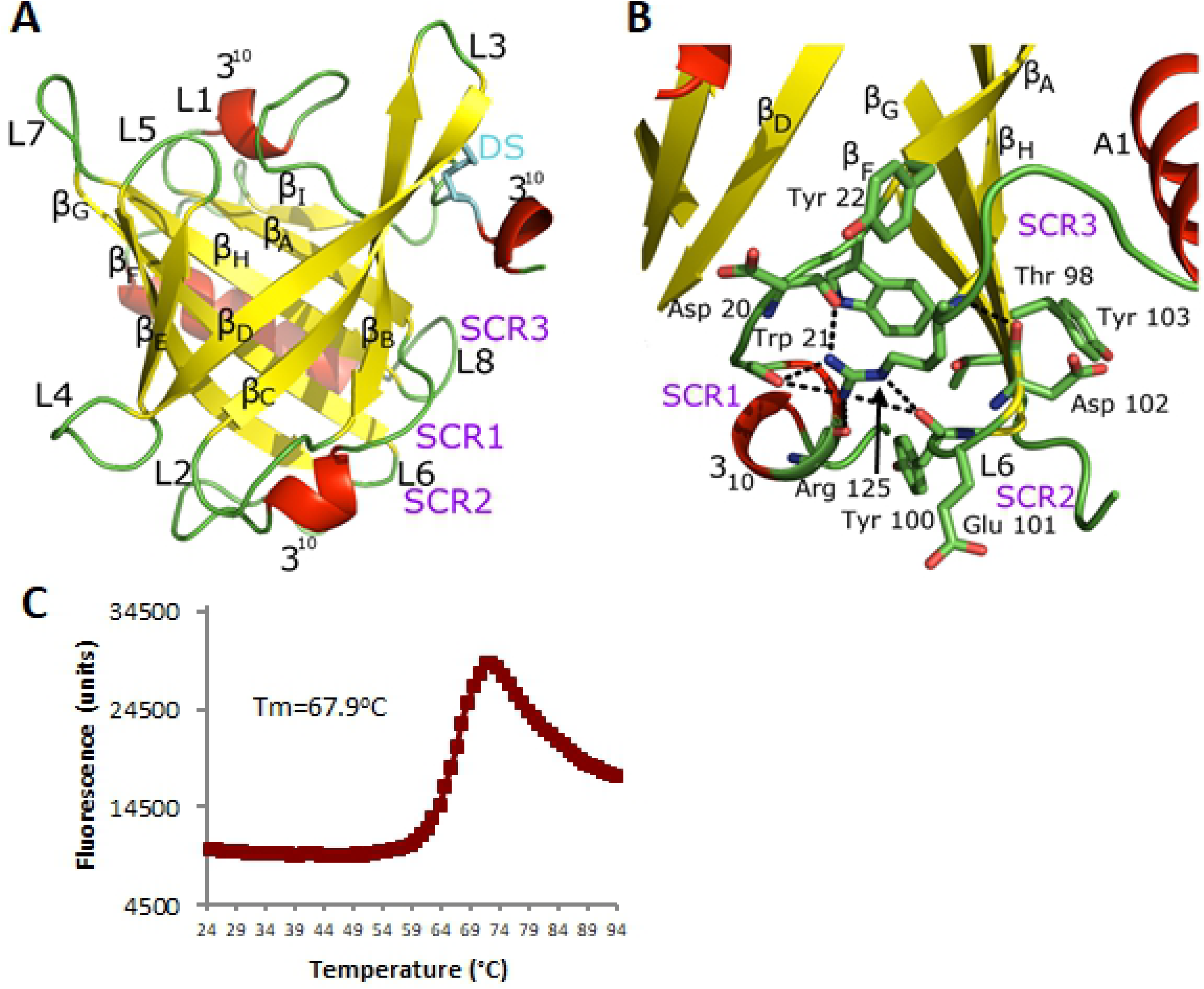
Comparison of predicted post translational modification sites. **(A)** Summary of glycosylation sites. **(B)** Three N-linked glycosylation residues (blue spheres) are predicted at Asn35, 52, and 75 in Can f 6; one in Can f 2 at Asn27; and two in Equ c 1 at Asn37 and 68. All these sites are on loops. The O-linked glycosylation site predicted in Can f 6 (purple spheres) is at the end of the C-terminus (Ser166) and is not modeled because this region of the structure is disordered. **(C)** Predicted phosphorylation sites (orange spheres). Phosphorylated Tyr22 and Ser18 in human tear lipocalin are present and labeled in Can f 6.

These differences in glycosylation and phosphorylation will affect lipocalin solubility, ability to bind other proteins and may also be an important feature of antibody recognition and lipocalin immunogenicity. Glycosylation and phosphorylation may mask or alter the surface features of nearby residues(39). For example, Mouse mAb 65 recognizes the natural glycosylated Equ c 1 and not the recombinant unglycosylated Equ c 1(23). Similarly, human tear lipocalins are known to be phosphorylated and changes in phosphorylation may play a role in lipocalin related diseases(38).

### Comparison of the ligand binding cavities of some lipocalins

In many cases the biologically relevant lipocalin ligands are as yet unknown. Thus close examination of the properties of their ligand binding cavities, deduced from the presence of unresolved electron density and the shape of the cavity volume in these locations, is of interest. In the structure of Can f 6 there is unmodeled electron density inside the highly hydrophobic ligand binding cavity and a uniquely shaped volume (Fig. 6A-C). Since Can f 6, as described here, was expressed in the bacterial periplasm and purified in its native form, the electron density observed in the ligand pocket may be from a full or partially occupied crystallization reagent, or metabolite from the bacteria used for expression. The ligand in the cavity is not thought to be from the crystallization process because the shape is inconsistent with the Poly Ethylene Glycol (PEG) used in the crystallization conditions. In either case the ligand molecule likely has similar properties to the natural binding ligand.

**Fig. 6.**
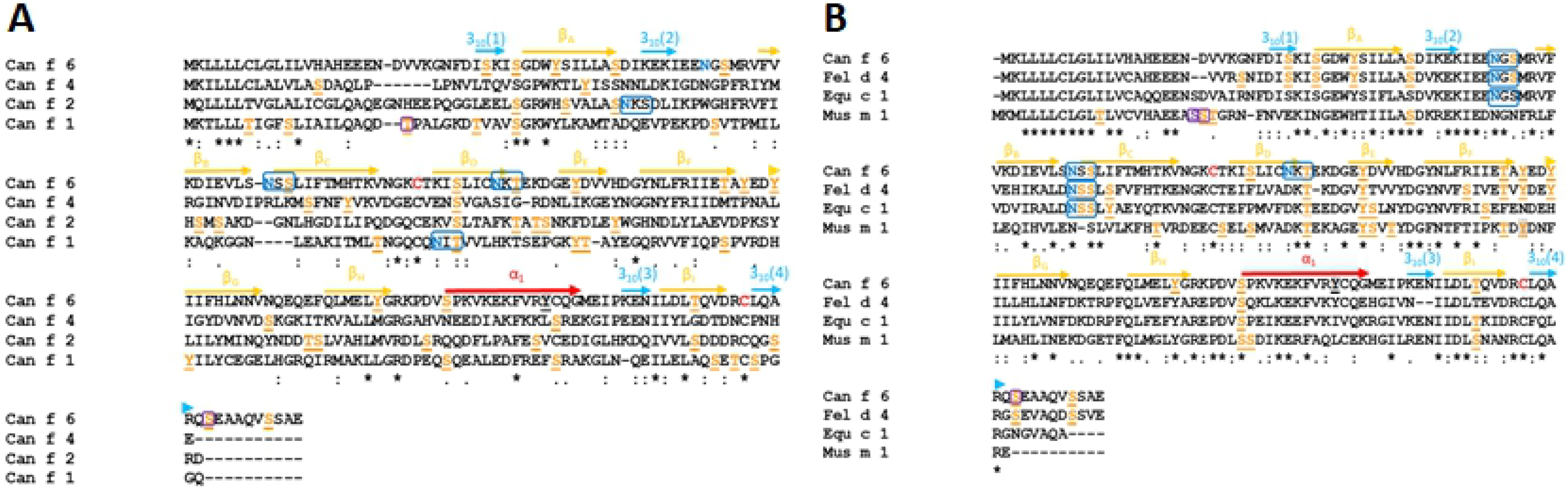
A comparison of the ligand binding cavities in the different lipocalins. Cavity residues and volume maps (red) were calculated using a rolling probe of 1.4Å and displayed as Connolly surfaces (25, 26, 27). **(A)** The overall ligand core is centrally located within the beta barrel and closed at the top by the L1 and L5 loops. **(B)** Residues from multiple strands of the β-barrel contribute to the ligand cavity core with hydrophobic amino acids and aromatic side chains in the interior of the ligand binding surface. **(C)** The ligand cavity volumes are fairly similar except for Equ c 1. **(D)** The ligand cavities of Can f 2 (red), Can f 4 (grey), Equ c 1 (purple), and Mus m 1 (brown) have a variable shape. Can f 6 and Equ c 1 have similar ligand cavity shape, with the slight differences between them likely reflects similarly shaped ligands.

The central cavities of the lipocalins described here vary in size and shape reflecting differences and likely specificity for their natural ligands (Fig. 6D). The cavity of Can f 6 has a solvent accessible surface area of approximately 132 Å^2^ with a volume of 55.3 Å^3^ calculated by CASTp (25). These sizes are comparable to that of Can f 2 which has a surface area of 152.0Å^2^ with a volume of 60.6Å^3^ and smaller than Can f 4 which has a surface area of 158.3 Å^2^ with a volume of 95.0Å^3.^ (Fig. 6C).

As observed for other lipocalins, the residues of Can f 6 ligand cavity are mainly hydrophobic. A total of 17 residues from the 8 antiparallel β-barrel are involved in defining the shape of the Can F 6 ligand cavity. Twelve of the 17 amino acids that line the cavity are hydrophobic with 4 polar and 1 negatively charged residue (Table 4). A comparison of the homologous lipocalins show little sequence conservation of the cavity except for a conserved Asn91 (Table 4) and highly conserved Val42, Phe93, and Leu108 (Fig. 6D)

**Table 4:** ligand binding pocket analysis of Can f 6 with the 17 main amino acid residues that define the shape of ligand binding cavity compared to homologous proteins.

| Can f 6 | 26L | 40M | 42V | 57F | 59M | 70L | 72L | 74C | 85V | 87H | 91N | 93F | 106F | 108L | 119L | 121E | 123Y |
| --- | --- | --- | --- | --- | --- | --- | --- | --- | --- | --- | --- | --- | --- | --- | --- | --- | --- |
| Can f 4 | Y | F | L | F | F | N | V | A | G | Y | N | F | Y | V | V | L | M |
| Can f 2 | A | F | V | G | L | V | L | A | L | Y | N | L | L | M | V | H | M |
| Equ c 1 | F | M | V | A | Y | F | M | F | L | Y | N | F | L | L | L | E | Y |
| Fel d 4 | L | M | V | F | H | L | L | A | V | Y | N | F | L | L | L | E | Y |

Only the amino acid position for Can f 6 is listed.

At the cavity opening of the lipocalins discussed here, L1 and L5 close the cavity entrance (Fig. 7). Unique to Can f 6 is His87 of L5 that closes the L1 loop over the ligand space via a hydrogen bond to the backbone carbonyl of Ser39 in L1. His87 is itself stabilized by intra L5 hydrogen bonds and L5 Asn91. The L1 lid loop is stabilized by numerous hydrogen bonds to itself and by L1 Arg41 binding to the backbone residues of the L1 3_10_ helix. In contrast, Can f 2, Fel d 4, Equ c 1, Mus m 1, and other lipocalins, have a highly conserved Tyr present instead of a His (Fig. 7). Though Tyr and His are similar, this site may be important for ligand recruitment and recognition. Nearby Asn91 and Arg41 are highly conserved within the lipocalins (Fig. 2) and alternate residues have strongly similar properties, indicating a highly conserved ligand calyx cavity capping mechanism for recognition and binding of ligands.

**Fig. 7.**
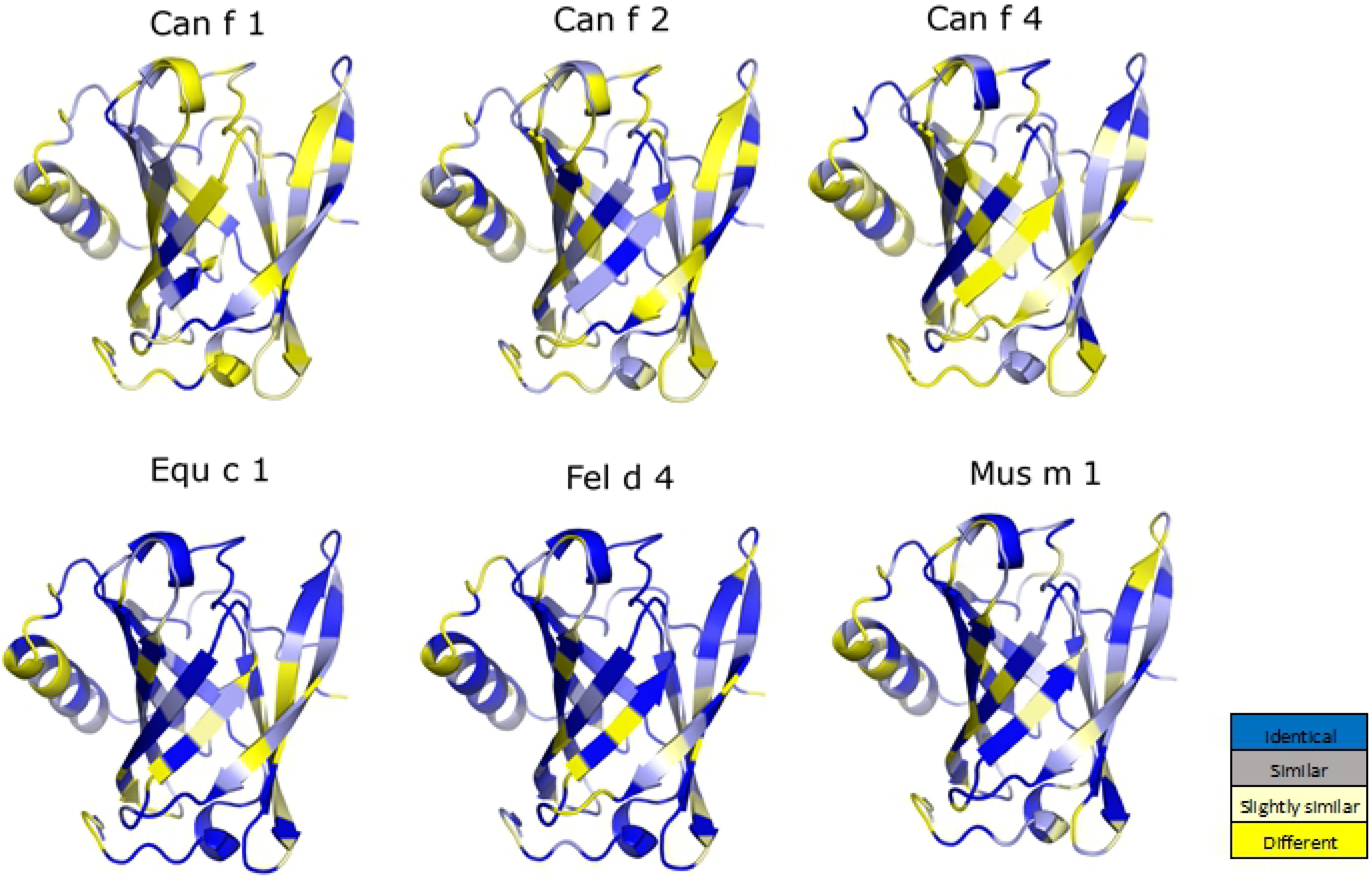
Comparison of the cavity capping residues of lipocalins. In Can f 6, at the entrance of the calyx cavity His87 in L5 stabilizes L1 over the ligand binding mouth. Arg41 and Asp91 stabilizes His87 and the L1 loop and are highly conserved indicating a common capping mechanism between lipocalins. The His in Can f 6 is unique compared to a highly conserved Tyr that is observed in the other lipocalins reflecting a unique ligand binding mechanism. Volume map is portrayed in red as Connolly surfaces (26).

The calyx cap conformation in Can f 6, places Met40 of the L1 loop over the ligand cavity where it does not make significant internal interactions. In the other lipocalins, an L1 Phe sits over the ligand space (Fig. 7) and these residues are thought to also be involved in ligand recognition and capture.

It is possible that the Can f 6 structure here represents a closed conformation with the unresolved internal electron density that could be a full or partial ligand. Other lipocalins have different conformations of L1 and L5. Compared to the structure of Can f 6, the structure of retinoic acid bound to RBP reveals that L1 and L5 are further apart(33) and a broader opened mouth conformation. The L1 and L5 conformation accommodates the sterol ring of the ligand while the aliphatic tail sits in the ligand cavity. However, the structure of the apo form of RBP does not have significant global changes in L1 and L5. By comparison, in the structure of the Sphingosine 1-Phosphate bound apolioprotein(34), the L1 and L5 are more similarly oriented to that observed for Can f 6 although the L1 loop is rearranged to a more open conformation. The resulting space is filled with waters that make hydrogen bonds to the sphingosine phosphate head group. Only a fully resolved ligand bound structure will reveal whether or not that is the case.

### Human IgE binds recombinant Can f 6

The ability of recombinant Can f 6 to bind human sera IgE was analyzed in 5 patients who were previously determined to be AP dog dander positive on immediate skin prick testing as well as dog dander IgE positive (>5 kU/L, median 33.4 kU/L, ImmunoCAP) (Table 5). Age matched controls who were immediate skin prick test negative and IgE negative (<0.35kU/L) were identified. The frequency of qualitative IgE binding to Can f 6 in dog dander positive samples was 60% (3 out of 5 patients, p-value=0.1199) (Fig 8). IgE reactivity verifies the antigencity of recombinant Can f 6.

**Table 5:**
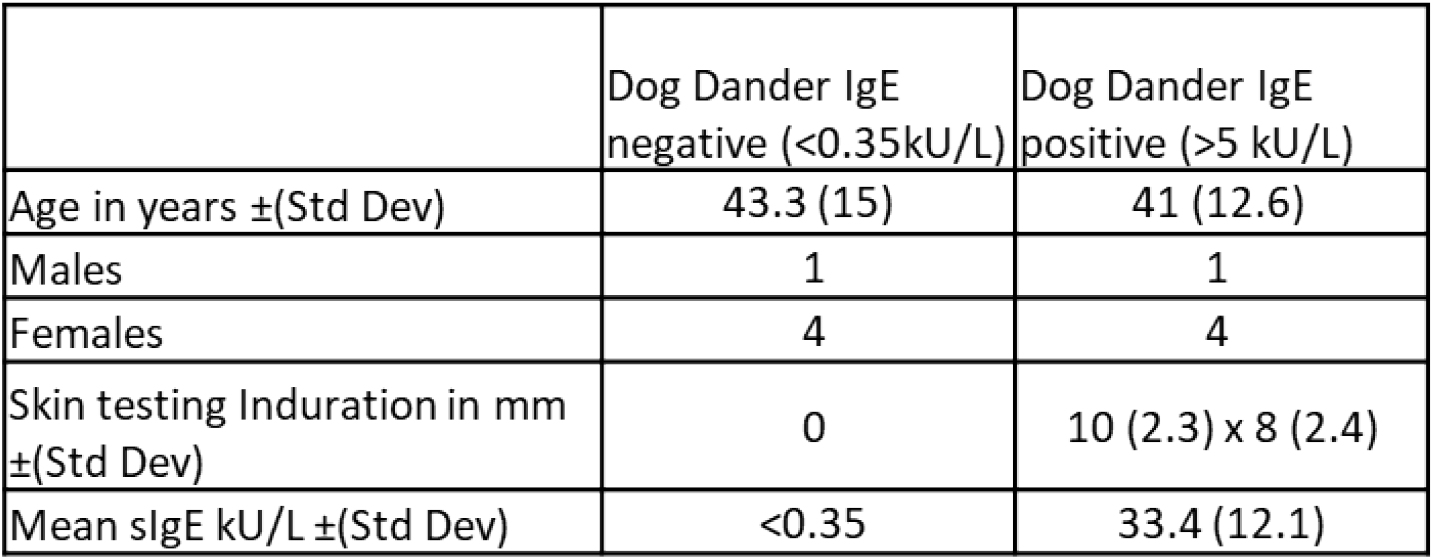
Demographics of Human Serum samples

|  | Dog Dander IgE<br>negative (<0.35kU/L) | Dog Dander IgE<br>positive (>5 kU/L) |
| --- | --- | --- |
| Age in years ±(Std Dev) | 43.3 (15) | 41 (12.6) |
| Males | 1 | 1 |
| Females | 4 | 4 |
| Skin testing Induration in mm<br>±(Std Dev) | 0 | 10 (2.3) x 8 (2.4) |
| Mean sIgE kU/L ±(Std Dev) | <0.35 | 33.4 (12.1) |

**Fig. 8:**
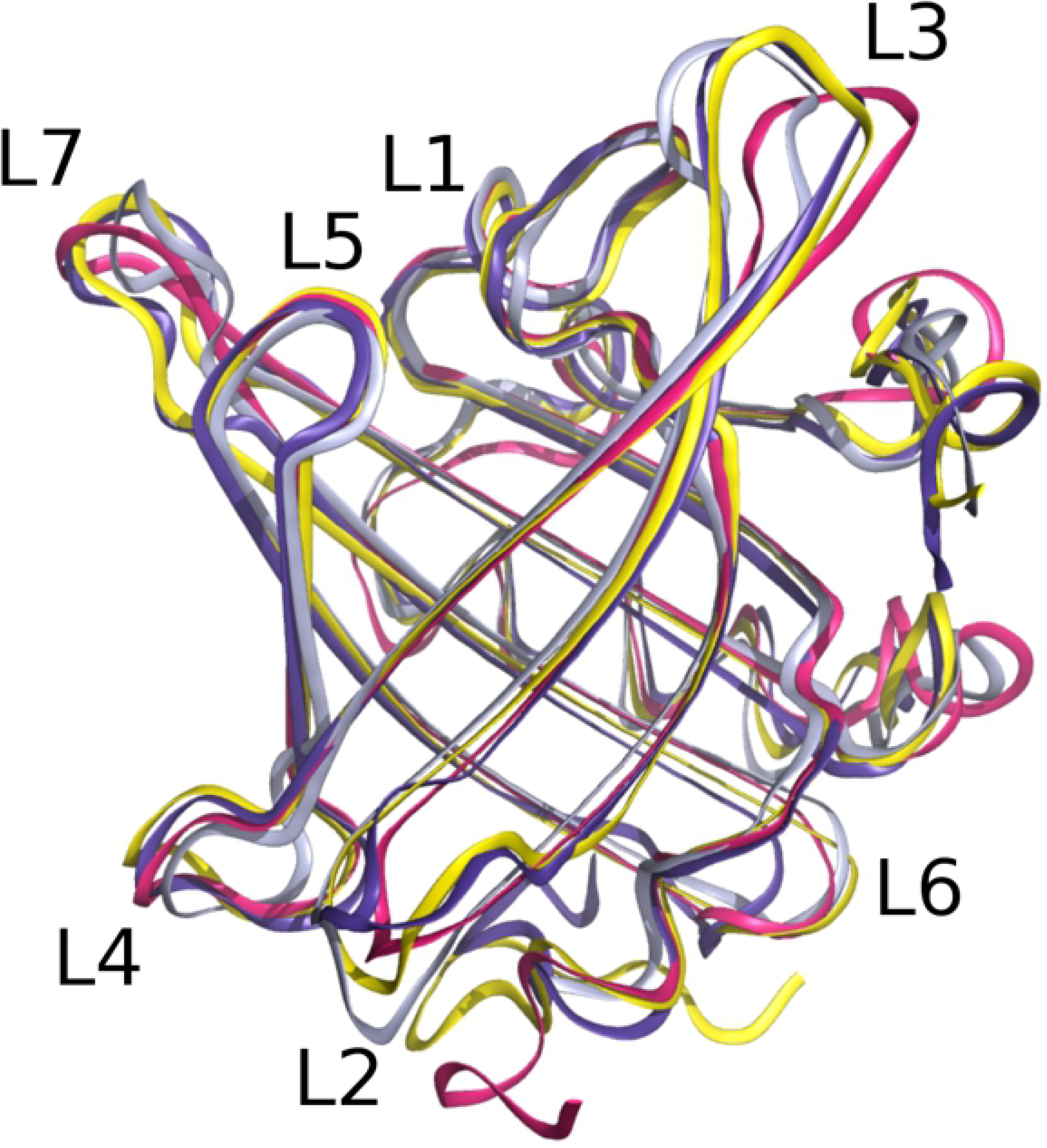
Highly pure recombinant Can f 6 binds human serum IgE in individuals who have been tested for sensitization to unfractionated whole dander IgE. Three of 5 samples from dog dander positive individuals have marked elevation in IgE binding to Can f 6. Mean and standard deviation of OD405nm for patients with dog IgE<0.35: 12.8±8.34 and IgE>5: 108±54.2 for Can f 6. Experiments were completed twice with the trend remaining the same. Overall p-value=0.1199.

Although no data was available regarding symptoms with exposure, molecular component resolved testing may be more accurate in the diagnosis of dog allergy especially in the individual who tested dog dander IgE and skin testing negative but who has evidence of IgE binding to Can f 6.

In summary, the biologically active high resolution x-ray structure of Can f 6 reveals greater insights into the specific roles of critical residues involved in ligand recognition and allergenicity. Further investigation into the role of lipocalins in vivo as well as in clinical allergenic diseases are needed.

## Acknowledgements

We acknowledge and thank the Advanced Photon Source at Argonne National Laboratory for the facilities and staff of the 24-ID-C beamline for their assistance in data-collection. We are grateful to Dr. Jennifer Maynard for the Mopac54 vector plasmid. Samples and data used for this study were downloaded from the National Jewish Health Research Database (https://rdb.njhealth.org/sec/r2prd/ep/home.cfm), supported by National Jewish Health.

